# PonA2’s contributions to biofilm and colony formation independent of its catalytic domains in *Mycobacterium smegmatis*

**DOI:** 10.64898/2026.06.03.729914

**Authors:** Bryan A Montero-Gutierrez, Jathin K Bande, Takehiro Kado

**Affiliations:** Department of Biology, Missouri State University, 901 South National Avenue, Springfield, MO 65897; Central High School, 423 E Central St, Springfield, MO 65802

**Author notes:** Correspondence, (TK).

**Keywords:** Mycobacteria, biofilm, colony, *ponA2*, penicillin-binding protein, membrane domain

## Abstract

Biofilm and colony growth creates microenvironments that require coordinated localization and function of cell surface molecules within the membrane. However, the mechanisms by which membrane organization regulates these surface molecules during these biofilm-associated growth are poorly understood, limiting our understanding of how bacteria adapt and survive in multicellular communities. *Mycobacterium smegmatis* contains an inner membrane domain (IMD) at the subpolar regions of the cell that helps mediate cell envelope synthesis. Prior research has identified *ponA2* as critical for de novo formation of distinct plasma membrane domains in planktonic growth. PonA2 is a penicillin-binding protein that catalyzes peptidoglycan synthesis by transglycosylase (TG) and transpeptidase (TP) activities. To investigate the role of PonA2 in membrane domain organization in biofilm and colony growth, wild-type, Δ*ponA2*, the complement strain (c*ponA2*), and catalytic inactive variants of PonA2 (TG-, TP-, and TG-/TP-) were analyzed. The IMD subpolar localization in wild-type was preserved in biofilm and colony growth, indicating that IMD localization is not exclusive to planktonic growth. Both biofilm and colony growth of Δ*ponA2* showed significant structural deformities compared to wild-type. In contrast, the catalytic inactive mutants produced biofilm and colony structures that resembled wild-type, suggesting that PonA2 has additional noncatalytic functions during multicellular growth. The IMD localization of the catalytic inactive mutants was minimally impacted, suggesting that neither catalytic domain is required for IMD localization in biofilm and colony growth. Together, these findings advance our understanding of the complex mycobacterial membrane biology.

**Importance:** Planktonic, pellicle biofilm, and colony growth expose mycobacteria to distinct environmental conditions that can affect cell-envelope organization and survival. Prior research has shown that mycobacteria form subpolar plasma membrane domains that support polar cell elongation in planktonic growth, but it remains unclear whether this organization is conserved in multicellular biofilm and colony growth, where cells experience nutrient and oxygen gradients and altered cell-to-cell interactions. Our current study analyzed the plasma membrane domain formation across the two growth states and showed that membrane domain localization is conserved, while the mechanisms required to maintain these domains differ depending on growth conditions. These findings suggest that mycobacteria use growth state specific mechanisms to coordinate membrane organization.

## Introduction

Members of the mycobacterium family possess a characteristic thick, waxy, and complex multi-layered cell envelope that makes them intrinsically resistant to many antibiotics (1, 2). The mycobacterial cell envelope is composed of five main layers: the capsule, the mycolic acid-rich mycomembrane, the arabinogalactan layer, the peptidoglycan sacculus, and the plasma membrane (3). The plasma membrane plays an integral role in cell envelope synthesis in mycobacterium species. *Mycobacterium smegmatis and Mycobacterium tuberculosis* partition its plasma membrane into two distinct regions: the inner membrane domain (IMD) and the conventional plasma membrane which is tightly associated with the cell wall (PM-CW) (4–7).

The IMD region is enriched with proteins and lipids involved in cell envelope biosynthesis and is proposed to synthesize precursors of cell envelope molecules (4, 5, 8). In contrast to the IMD, the PM-CW is proposed to assemble cell envelope precursors synthesized in IMD into the mature cell envelope (4). The spatial localization of IMD also suggests its importance in cell elongation. The IMD is localized in the subpolar region of a growing *M. smegmatis* and *M. tuberculosis* (4–6). The elongation process of *M. smegmatis* differs from that of *Escherichia coli* in that *M. smegmatis* elongates from its poles rather than along the lateral cell body (9). Additionally, *M. smegmatis* grows asymmetrically at its poles as the old pole grows faster than the new pole (9). This results in daughter cells that have different growth rates, sizes, and distribution of macromolecules, which has been shown to aid in antibiotic immunity (10). The biosynthesis of de novo mycobacterial cell envelope requires coordinated and localized protein machinery at the cell poles to mediate the growth.

Previous studies have identified *ponA2* as a candidate gene involved in membrane domain formation (11). PonA2 is a nonessential class A penicillin-binding protein with bifunctional transglycosylase and transpeptidase activity (12). Biochemical fractionation experiments showed that PonA2 is primarily associated with the PM-CW (4). Deletion of *ponA2* caused delayed membrane domain formations, suggesting that PonA2 contributes to spatial coordination of envelope biosynthesis (11).

Those previous studies in membrane domain formations, however, have only been explored in planktonic growth (4, 5, 8, 11, 13–20), but its role in other multicellular growth states have not been explored. Cells in planktonic growth experience a relatively uniform distribution of nutrients, but colony and biofilm growth generate nutrient gradients that produce heterogeneous subpopulations (21). Since mycobacterial cell surface molecules play an important role in pellicle biofilm formation by promoting hydrophobic intercellular interactions (22), proper spatial organization of cell envelope biosynthesis could be important for generating the surface properties required for colony and biofilm formation. Therefore, proteins involved in lateral membrane partitioning, such as PonA2, may play a role in coordinating envelope organization.

Here, we strive to address the functionality of PonA2 in membrane domain organization in non-planktonic growth. We observed the insignificance of PonA2’s catalytic domains in macroscopic architecture formation, IMD localization, and cellular morphology in both biofilm and colony growth states. In contrast, the deletion of *ponA2* revealed the importance in biofilm and colony formation, as Δ*ponA2* produced smooth macroscopic architecture, suggesting the presence of an additional non-catalytic function of PonA2.

## Materials and Methods

### Cell Growth

For planktonic growth, cells obtained from the frozen stock of each *M. smegmatis* mc^2^155 strain was cultured in Middlebrook 7H9 supplemented with 15 mM NaCl, 0.2% (w/v) glucose, and 0.05% (v/v) Tween-80, and incubated at 37 °C with shaking at 160 rpm for 2 days. For isolated colony growth, 20 µL of planktonic stationary growth phase culture was inoculated in fresh 20 mL Middlebrook 7H9 medium supplemented with 11 mM glucose, 14.5 mM NaCl, and 0.05% (vol/vol) Tween 80. The liquid culture was incubated at 37 °C with shaking at 160 rpm for 16 h. Then 5 µL of the culture was dropped onto a Middlebrook 7H10 agar plate that was supplemented with 15 mM NaCl and 0.2% (w/v) glucose. The Middlebrook 7H10 agar plates were placed in a sealed plastic bag to prevent excessive dehydration and incubated at 37 °C for 4-6 days. Colonies were photographed with an iPhone 15 Pro Max camera (Apple). For pellicle biofilm growth, 20 µL of planktonic stationary growth phase culture was inoculated into 2 mL M63 media in a 24-well plate. The outer wells were filled with 2 mL of sterile water to minimize evaporation. The 24-well plates were incubated at 37 °C with no shaking for 5-7 days. Biofilms were photographed with the One Plus Nord N200 camera (BBK Electronics).

### Microscopy and quantification of IMD localization and cell morphology

Five microliters of *M. smegmatis* cells expressing HA-mCherry-GlfT2 were placed onto a 2% agar pad made of PBS. All images were taken using a Leica CTR6500 microscope (Leica Microsystems) equipped with a Hamamatsu C4742-95 camera and a Leica HCX PL APO 100×/1.40 OIL PH3 objective. The images were all set to have 2-ms phase exposure and 360-ms TXRed exposure. Microscopy images were processed using ImageJ (Fiji), and cells were outlined and segmented using Oufti (23). Fluorescence signals of each cell were analyzed using custom-written MATLAB codes that quantified the fluorescence intensity (24). MATLAB output was then graphed in Excel (Microsoft).

For analyzing cell lengths, we gathered the arbitrary length of each cell from the output file from Oufti and divided the arbitrary cell lengths by 10.6 pixel/μm to get the real cell length for all cells. We then took those cell lengths and used SuperPlotsofData (25) to generate the super plot for cell length. All experiments were repeated three times independently of each other.

Colony diameters were measured using Image J (Fiji) by drawing a straight line across the widest part of each colony. Pixel-based measurements were calibrated using the known diameter of the agar plate to calculate the true colony diameter. Colony diameters were analyzed using GraphPad Prism to generate the super plot for colony diameter. All experiments were repeated three or six times independently of each other.

### Cell Wall Staining

To visualize peptidoglycan synthesis in *M. smegmatis* colony cells, the scraped cells from an agar plate were placed in 1 mM RADA and incubated for 1 h at room temperature. Following incubation, cells were harvested by centrifugation at 3,000 rpm for 3 min, washed 3 times with filtered phosphate-buffered saline (PBS), fixed in 4% paraformaldehyde for 16 h at 4 °C, followed by 2 more PBS washes.

RADA incorporation was quantified using fluorescence microscopy images. For each field of view, the total number of cells were counted using the phase-contrast image. The TxRed image of the same field of view was used to count the number of cells that showed detectable RADA fluorescence. The percentage of RADA positive population was calculated by dividing the number of fluorescent cells by the total number of cells in the phase-contrast image and multiplied by 100. We then took the RADA positive population percentage values and used SuperplotsofData (25) to generate the super plot to display the percentage of RADA positive cells in each sample. Three field of views were analyzed for each biologically independent experiment.

## Results

### Subpolar localization of IMD was observed in both colony and biofilm growth of *M. smegmatis*

To analyze whether the IMD was formed in both colony and biofilm growth conditions in *M. smegmatis*, we gathered cells from both colony and biofilm structures and observed them under a fluorescence microscope to detect for a fluorescently tagged IMD-associated protein, galactofuranosyltransferase (GlfT2) (5). The macroscopic inspection of the structures revealed that *M. smegmatis* formed rugose colonies and robust pellicle biofilms (Fig. 1A). A microscopic examination revealed that both growth conditions retained their bacillary morphology and exhibited an increased fluorescence intensity of the IMD marker GlfT2 at their subpolar region (Fig. 1B). Quantitative analysis of the IMD fluorescence confirmed that the fluorescence intensity peaked in subpolar regions of the cell, with reduced fluorescence intensity in the mid-cellular regions (Fig. 1C). These trends were observed in both growth conditions with minor differences in fluorescence intensities. Overall, the IMD was formed at the subpolar region of the cell in both colony and biofilm growth states.

**Figure 1.**
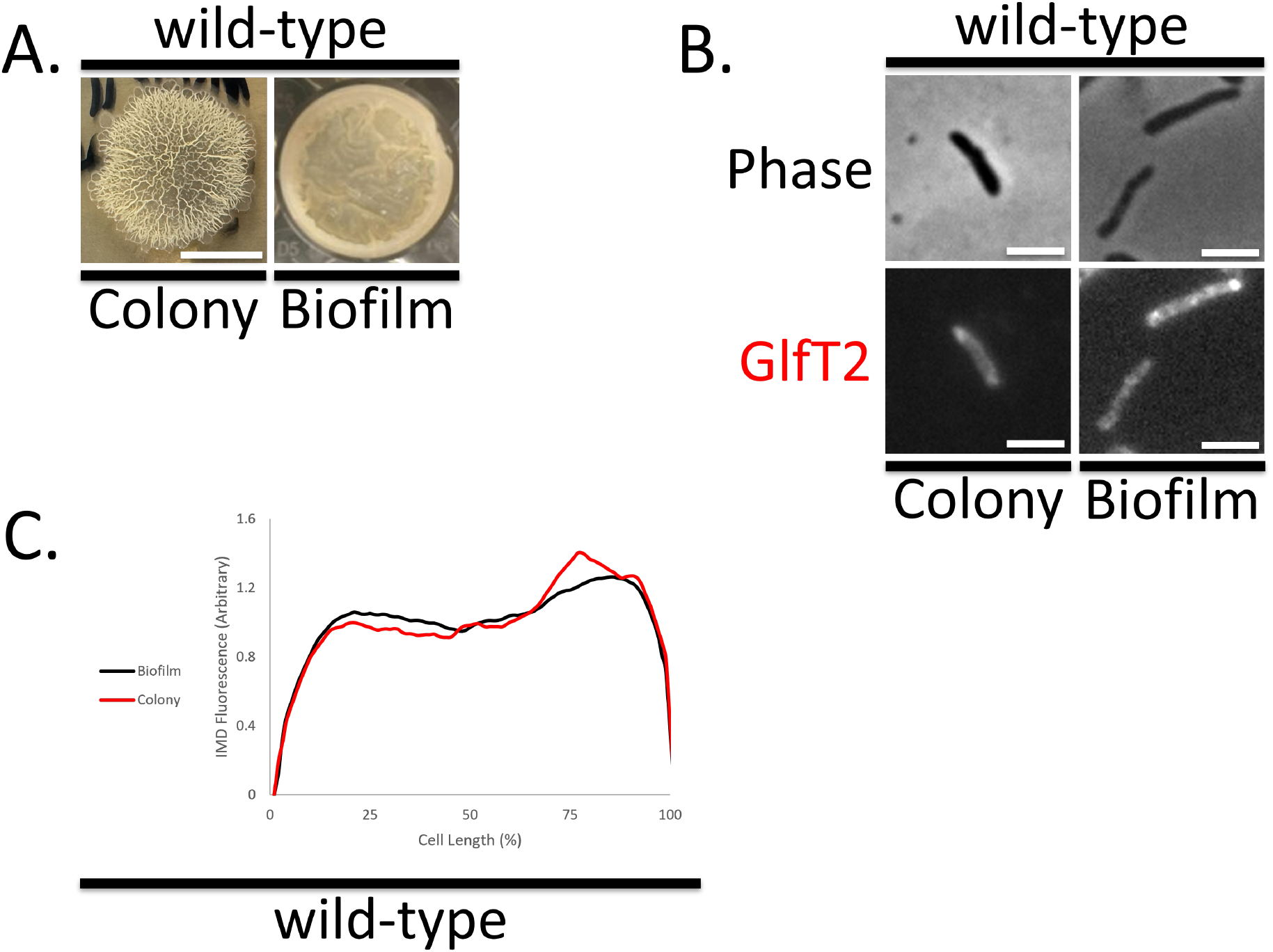
Subpolar IMD localization was observed in colony and biofilm growth of *M. smegmatis*. **A**. Colony growth was attained by spotting 5 µL of planktonic culture onto a 7H10 Middlebrook agar plate supplemented with DC and incubated at 37 °C for 5-7 days. Scale bar = 1 cm. Biofilm growth was attained by the inoculating 2 mL M63 media with 20 µL planktonic culture and was incubated at 37 °C for 5-7 days. **B**. Representative images of both growth conditions. Phase-contrast microscopy shows cell morphology while GlfT2 marks IMD localization. Scale bar = 2.5 µm. **C**. Data represents the average fluorescence intensities of GlfT2 as they occur along the entirety of the cell for colony (black) and biofilm (red) cells. Data was gathered from three biologically independent experiments, n > 230 cells for colony and biofilm conditions.

### The deletion of *ponA2* altered colony structure, biofilm structure, and cell morphology in colony growth

To determine the role of PonA2, which helps de novo IMD formation (11), in colony and biofilm growth, we compared colony morphology, biofilm formation, and IMD localization between wild-type and Δ*ponA2*. A macroscopic examination of Δ*ponA2* displayed significant structural differences in colony and biofilm structures (Fig. 2A). Δ*ponA2* formed smaller, smoother colonies and smooth-surfaced pellicle biofilms, deviating from the structured, organized colonies and pellicle biofilms that wild-type produced (Fig. 2A). Shifting to a microscopic scale, Δ*ponA2* cells in colony growth showed a shorter rod morphology, whereas Δ*ponA2* cells maintained similar cellular morphology and IMD localization in the biofilm growth compared to wild-type cells (Fig. 2B, C). In addition, there was an absence of IMD marker in Δ*ponA2* cells in the colony growth (Fig. 2B). The macroscopic morphological differences observed in both growth conditions between Δ*ponA2* and wild-type suggest that PonA2 plays an important role in structural growth process of both colony and biofilm structures (Fig. 2A). The microscopic morphological differences observed between the Δ*ponA2* growth conditions suggest that PonA2 plays a larger role in colony growth than biofilm growth. In colony growth, PonA2 was needed to keep the bacillary morphology and organize IMD at the subpolar region; however, PonA2 was not needed in biofilm growth to maintain bacillary morphology and IMD subpolar localization (Fig. 2B). Quantitative analysis of the GlfT2 fluorescence revealed that IMD is still localized in the subpolar regions of biofilm growth in both strains (Fig. 2C). These findings indicate that PonA2 is required for proper colony formation, biofilm formation, and IMD organization in colony growth.

**Figure 2.**
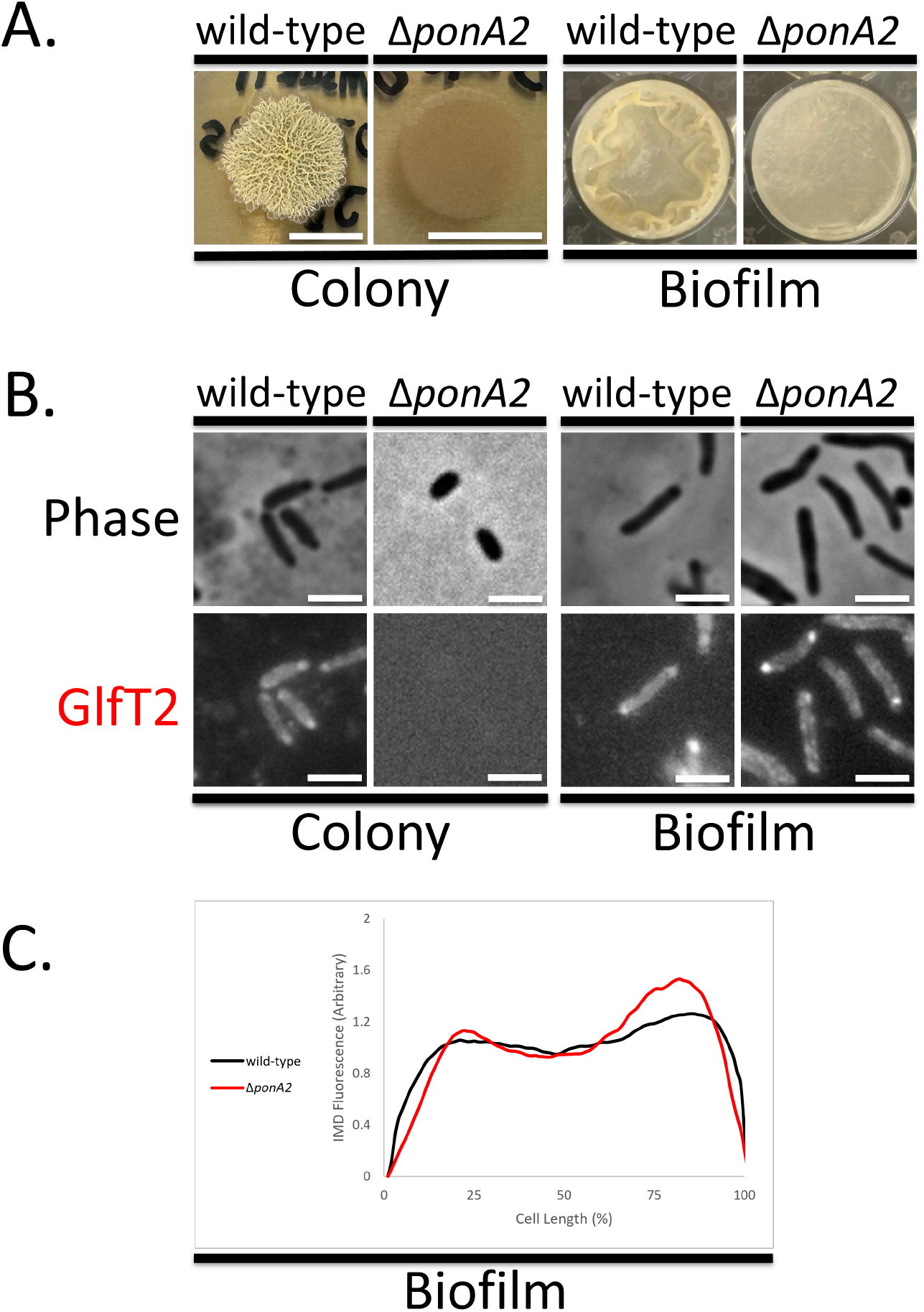
Comparative analysis of colony and biofilm structures, cell morphology, and IMD localization between Δ*ponA2* and wild-type. **A**. Colony growth was analyzed by plating 5 µL planktonic culture to a 7H10 Middlebrook plate supplemented with DC and incubated 37 °C for 5-7 days. Scale bar = 1 cm. Biofilm growth was procured by inoculating 2 mL M63 media with 20 µL planktonic culture. Incubation time was for 5-7 days at 37 °C. **B**. Representative images of both strains in both growth conditions. Scale bar = 2.5 µm. **C**. Data represents the average fluorescence intensities of GlfT2 as they occur along the entirety of the cell for wild-type (black) and Δ*ponA2* (red) cells in biofilm conditions, n > 230 cells for colony and biofilm conditions.

### The catalytic domains of PonA2 played a minor role in colony and biofilm growth

PonA2’s classification as a class A-penicillin binding protein means it contains both transglycosylase (TG) and transpeptidase (TP) catalytic domains (11, 26). To determine whether the catalytic domains of PonA2 are necessary for colony and biofilm growth, we compared the colony and biofilm morphology by using catalytically inactive mutants, that were generated in the previous study (11). The Δ*ponA2* strain showed less rugose and more uniform morphology in both colony and biofilm (Fig. 2A, 3A). In contrast, the complimented strain restored both colony and biofilm morphology to wild-type appearances (Fig. 3A). Both of the single catalytic mutant strains (TG-, TP-) and the double catalytic mutant strain (TG-/TP-) had similar colony and biofilm appearance to wild-type (Fig. 3A) while TG-/TP- showed significantly increased colony diameter (Fig. 3B). These results demonstrate that the catalytic domain of PonA2 played a minor role in rough colony and pellicle biofilm formation, suggesting that PonA2 contributes to these processes through non-catalytic function.

**Fig. 3.**
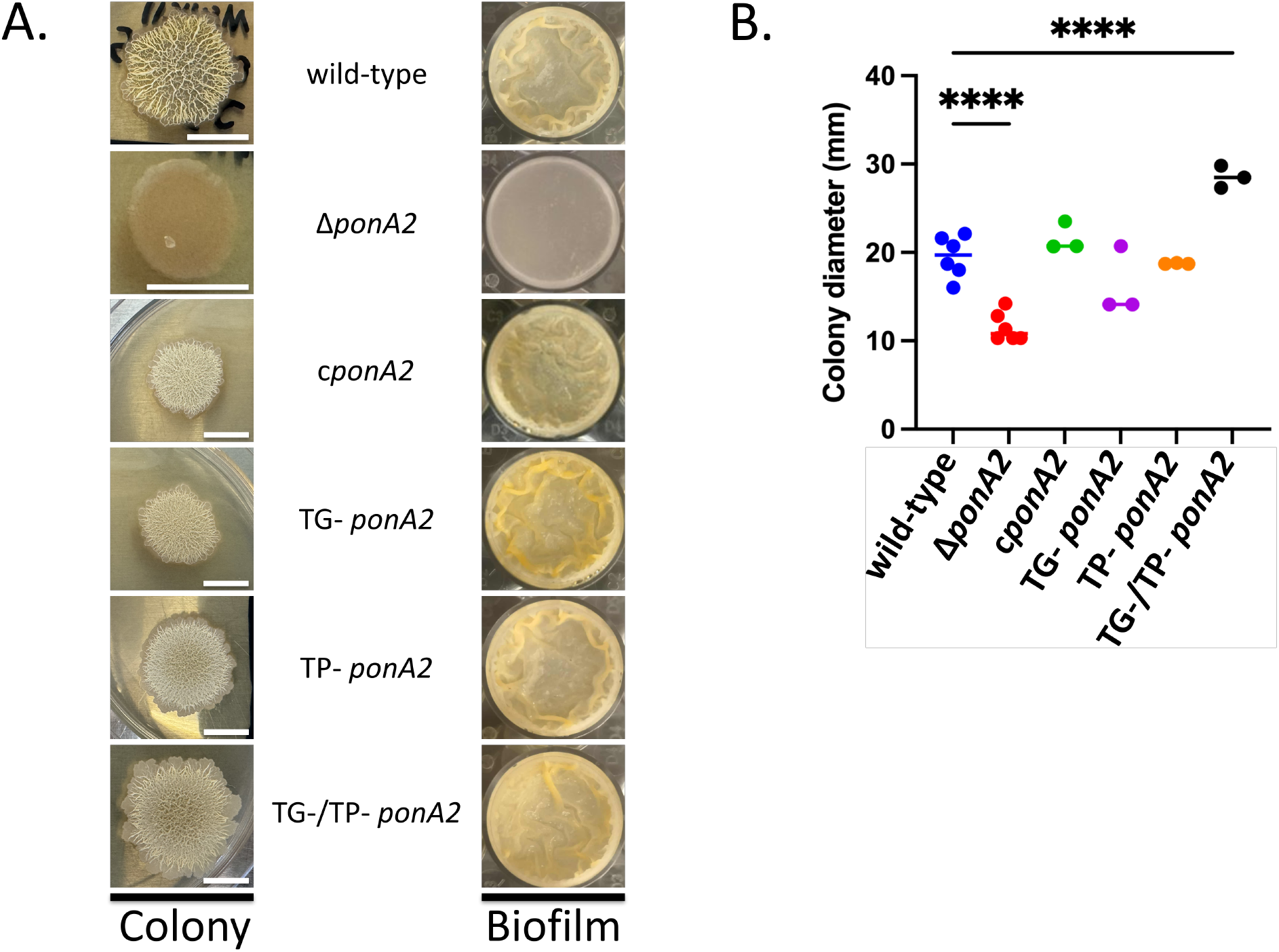
Macroscopic structural comparison of catalytic mutant strains to wild-type. **A**. Colony growth was established by spotting 5 µL of planktonic culture onto a 7H10 Middlebrook plate supplemented with DC and incubated at 37 °C for 6 days. Scale bar = 1 cm. **B**. Colony diameter measurements for the various strains. Each point represents an individual colony. Statistical significance was determined by the Kruskal–Wallis test, followed by Dunn’s multiple-comparison test. ns, no statistically significant difference; ^****^p<0.001. Data was obtained from three or six biologically independent experiments.

### The catalytic domains of PonA2 played a minor role in IMD localization and cell length

To determine whether the catalytic domains of PonA2 play a role in IMD localization in both colony and biofilm, IMD localization of the mutant strains were analyzed. The wild-type strain showed bacillary morphology and subpolar IMD localization in both colony and biofilm growth conditions (Fig. 1AB, 4AB). Δ*ponA2* exhibited shorter rod morphology and no IMD localization in colony growth but showed normal rod morphology and polar IMD localization in biofilm growth conditions (Fig. 2BC, 4AB). The compliment strain, TG-, TP-, and TG-/TP- strains all exhibited bacillary morphology and subpolar IMD localization in both colony and biofilm growth conditions (Fig. 4AB). Analysis of cell length revealed that Δ*ponA2* cells in colony growth were significantly shorter compared to wild-type, whereas the complimented and catalytic mutant strains were comparable to wild-type (Fig. 4C). In biofilm growth, no significant differences in cell length were observed between wild-type and the rest of the strains (Fig. 4C). These findings indicate that the inhibition of the catalytic domains of PonA2 does not impair IMD organization, suggesting that PonA2 has an additional unknown function in addition to its catalytic domains to form the plasma membrane domain.

**Figure 4.**
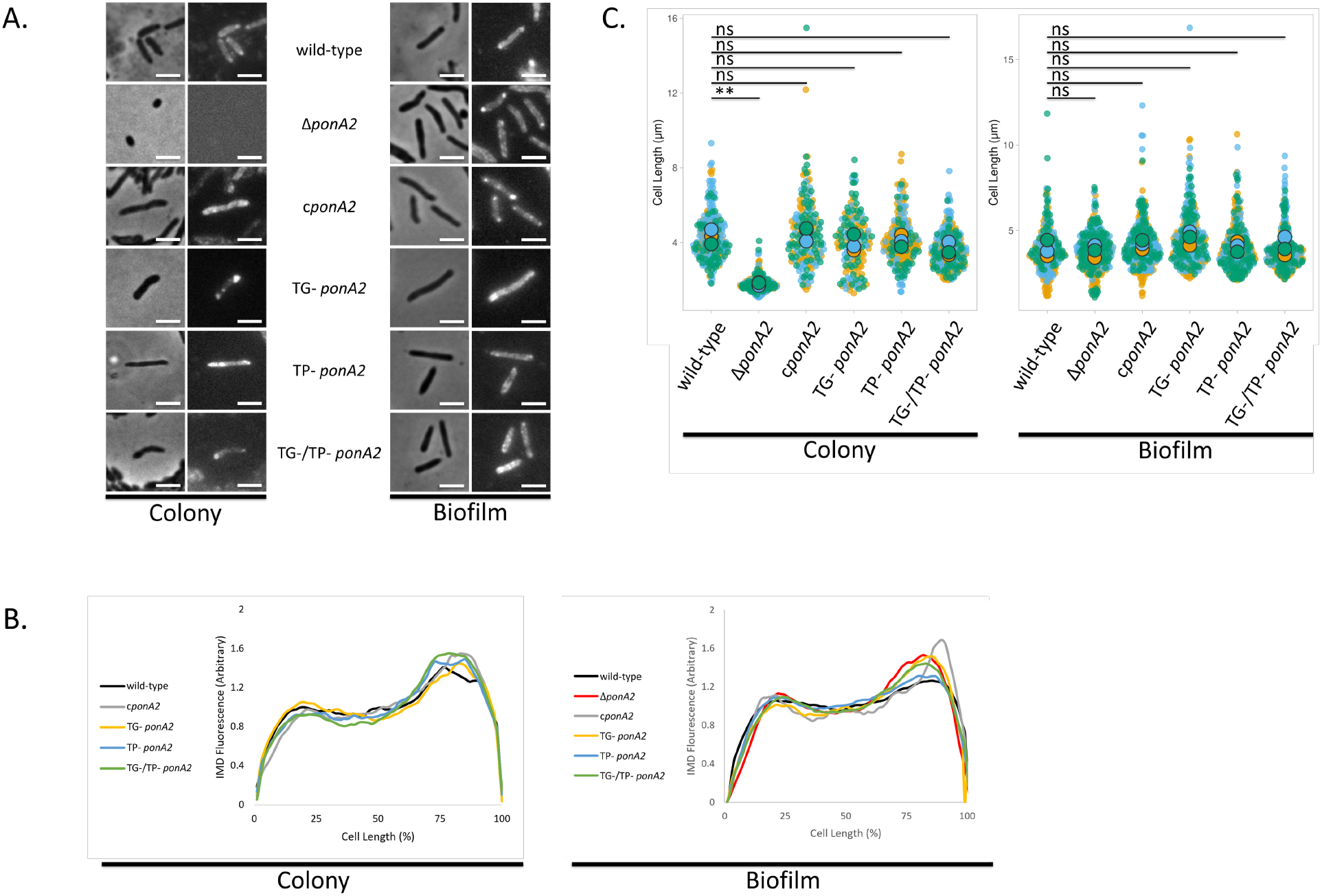
Microscopic analysis of cell morphology, IMD localization, and cell length. **A**. Representative images of all catalytic mutant strains in both growth conditions. Scale bar = 2.5 µm. **B**. Data represents the average fluorescence intensities (n > 230 cells) of GlfT2 as they occur along the entirety of the cell for each catalytic mutant strain. **C**. Cell length analysis of the multiple strains in both colony and biofilm growth. Each color in each condition represents one experiment, with the larger circles representing the average of the respective experiment. Statistical significance was determined using Welch’s t-test. ns, no statistically significant difference; ^**^p<0.01. Data was obtained from three biologically independent experiments.

### Deletion of *ponA2* showed ongoing cell wall synthesis in colony growth

To determine whether PonA2 affects ongoing cell wall synthesis, we used the fluorescent D-amino acid probe RADA (27, 28) in wild-type and Δ*ponA2* cells in colony growth conditions. Since Δ*ponA2* cells displayed shortened morphology and loss of subpolar IMD localization, we hypothesized that deletion of *ponA2* would reduce peptidoglycan synthesis in colony growth. The RADA successfully incorporated in both wild-type and Δ*ponA2* cells indicating that cell wall synthesis continues in the absence of *ponA2* (Fig. 5A). Quantitative analysis of RADA incorporation revealed no significant difference in RADA incorporation between wild-type and Δ*ponA2* in colony growth (Fig. 5B). These findings indicate that the deletion of *ponA2* shortened cell wall without inhibition of cell wall synthesis in colony growth.

**Figure 5.**
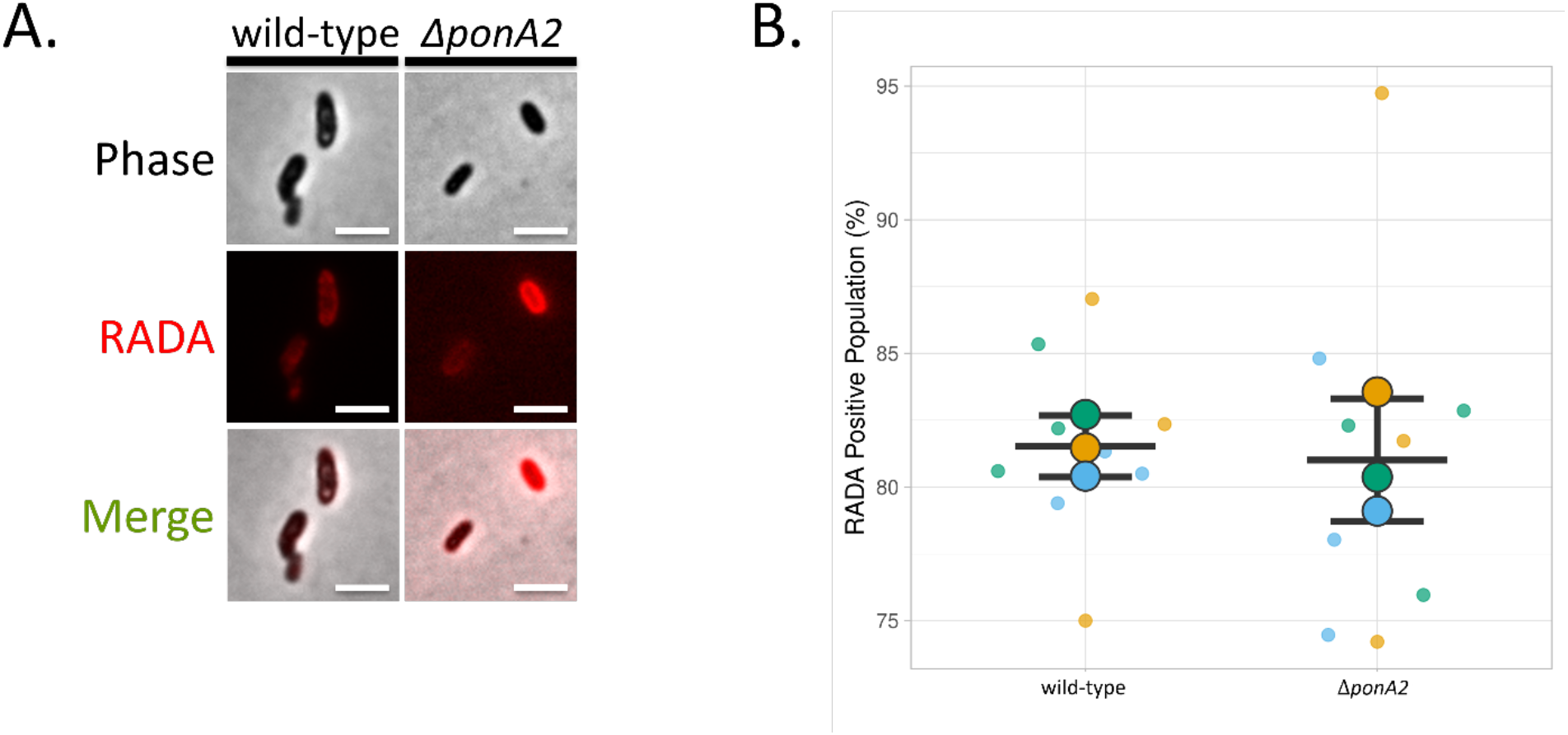
RADA incorporation of both wild-type and ΔponA2 in colony growth conditions. **A**. Cells scraped from colony culture were exposed to 1 mM RADA for 1 h, followed by three PBS washes, then exposed to 4% paraformaldehyde for 30 min, followed by two final PBS washes. **B**. Displays the percentage of cells that incorporated RADA into their cell in each condition. Each color in each condition represents one experiment, with the larger circles representing the mean of the experiment. The longer horizontal line represents the mean the three experiments, and the smaller lines represents the standard deviation. Statistical significance was assessed using Welch’s t-test using data obtained from three biologically independent experiments, with the sample size of each datum ranging from 56 < n < 274. No statistical significance was observed between the two conditions (p = 0.75).

## Discussion

Previous models of PonA2 have traditionally characterized the protein down to its transglucosylase and transpeptidase catalytic functions; however, the findings from this study challenge this model by suggesting that PonA2 may also possess non-catalytic functions. This claim is supported by the observation that the catalytic inactive variants of PonA2 maintained colony and pellicle biofilm architecture, bacillary morphology, and subpolar IMD localization comparable to wild-type. In contrast, the deletion of *ponA2* produced abnormally smooth and smooth-surfaced colonies and pellicle biofilms. In colony growth conditions, Δ*ponA2* cells displayed a shortened bacillary morphology and the loss of a detectable IMD localization, whereas in pellicle biofilm growth conditions, the deletion of *ponA2* did not impact cell morphology and IMD localization. From these data, we concluded that PonA2, but not its catalytic domains, is necessary for normal colony and pellicle formation.

The mechanism of how surface alterations resulting from the *ponA2* deletion affect the macromolecular architecture of colonies and biofilms is still unclear. We propose that PonA2 facilitates the proper levels of cell-to-cell adhesion required to create the complex folds and ridges seen in colony and biofilm structures. Cell-to-cell adhesion within structured communities is thought to mainly be attributed to the hydrophobic nature of the mycobacterial cell envelope (29). The mycobacterial cell envelope is composed of up to 60% of lipids, which is what gives the mycobacterial cell its strong hydrophobic tendency (30, 31). We postulate that the deletion of *ponA2* may indirectly, through some unknown mechanism, decreases the total lipid composition of the mycobacterial cell envelope, and therefore decrease its hydrophobic force and intercellular adhesion abilities. Following this logic, a reduction in hydrophobicity would decrease the cording ability of the mycobacteria, which may explain the smooth topology observed in the macrostructure of Δ*ponA2* colony and biofilm.

Experiments with Δ*ponA2* cells during colony growth demonstrated IMD-independent cell wall synthesis and reduced cell length. The shorter length was also identified in point mutants of DivIVA, a cytoskeletal protein in mycobacteria (6, 32). The point mutation experiments demonstrated altered peptidoglycan synthesis and subtle IMD delocalization in planktonic growth (6, 32). While the pulldown did not show direct interaction between DivIVA and PonA2 (33), both proteins associate with the PM-CW in *M. smegmatis* (5, 17). PonA2 and DivIVA might interrelate to manage the cell elongation during non-planktonic growth.

The functions of PonA2 beyond its catalytic activity remain unclear. Previous research with *M. tuberculosis* identified 6 genes essential for both colony and pellicle biofilm growth (34). The 6 genes include *ponA2, glpK, fadA2, mmaA4, cysQ*, and Rv3779. Of these, *mmaA4* is of interest as it encodes for mycolic acid methyltransferase, an enzyme that synthesizes oxygenated mycolic acids in the mycomembrane (35, 36). Although *M. smegmatis* does not contain the *mmaA4* gene, it does contain several related mycolic acid methyltransferase-like enzymes that modify *M. smegmatis* mycolic acids (37). Disruptions in mycolic acid-modifying enzymes, such as MSMEG_0913, have been shown to alter specific *M. smegmatis* mycolates, supporting the broader idea that mycolic acid modifying enzymes can influence cell envelope composition (38). These facts provide a possible bridge to our aforementioned hydrophobicity model; if the deletion of *ponA2* indirectly alters the organization or activity of mycolic acid-modifying enzymes, the resulting changes in envelope lipid composition could reduce the overall hydrophobicity of the cell, reducing the possibility of cell cording and contribute to the smooth colony and biofilm topology observed in Δ*ponA2*. Therefore, PonA2 may have a non-catalytic function in coordinating peptidoglycan synthesis and lipid-modifying pathways required to maintain cell envelope architecture.

While further studies examining in interactions between PonA2, -DivIVA, and -mycolic-acid-methyltransferases, as well as cell envelope composition and hydrophobicity of *ponA2* deletion mutant will be needed to fully elucidate the non-catalytic function of PonA2, we have shown that the physical presence of PonA2 is important for biofilm and colony growth. Our findings expand the current model of PonA2 by suggesting that its contribution to mycobacterial growth extend beyond its established transglycosylase and transpeptidase activities.

## Acknowledgement

We thank Dr. Yasu S. Morita and Dr. M. Sloan Siegrist at University of Massachusetts Amherst for support related to material transfer and strain sharing. We are also grateful to Dr. Kyoungtae Kim at Missouri State University for sharing GraphPad Prism.

